# The role of four cholesterol-recognition motifs localized between amino acid residues 400-550 in regulating translocation and lytic activity of Adenylate Cyclase Toxin

**DOI:** 10.1101/2021.11.26.470122

**Authors:** Jone Amuategi, Rocío Alonso, Helena Ostolaza

**Author notes:** Corresponding author: Helena Ostolaza,. Biofisika Institute (UPV/EHU, CSIC) and Department of Biochemistry and Molecular Biology, University of Basque Country (UPV/EHU), 48080 Bilbao, Spain.

## Abstract

Adenylate Cyclase Toxin (ACT or CyaA) is an important virulence factor secreted by *Bordetella pertussis*, the bacterium causative of whooping cough, playing an essential role in the establishment of infection in the respiratory tract. ACT is a pore-forming cytolysin belonging to the RTX (Repeats in ToXin) family of leukotoxins, capable of permeabilizing several cell types and pure lipid vesicles. Besides, the toxin delivers its N-terminal adenylate cyclase domain into the target cytosol, where catalyzes the conversion of ATP into cAMP, which affects cell signalling. In this study we have made two major observations. First, we show that ACT binds free cholesterol, and identify in its sequence 38 potential cholesterol-recognition motifs. Second, we reveal that four of those motifs are real, functional cholesterol-binding sites. Mutations of the central phenylalanine residues in said motifs have an important impact on the ACT lytic and translocation activities, suggesting their direct intervention in cholesterol recognition and toxin functionality. From our data a likely transmembrane topology can be inferred for the ACT helices constituting the translocation and the hydrophobic regions. From this topology a simple and plausible mechanism emerges by which ACT could translocate its AC domain into target cells, challenging previous views in the field. Blocking the ACT-cholesterol interactions might thus be an effective approach for inhibiting ACT toxicity on cells, and this could help in mitigating the severity of pertussis disease in humans.

## INTRODUCTION

*Bordetella pertussis* causes in humans a highly contagious respiratory infection known as whooping cough (pertussis), which remains a significant cause of disease and death in infants worldwide [1–3]. The bacterium produces several virulence factors, among which the **A**denylate **C**yclase **T**oxin (ACT or CyaA) is crucial for colonization of the respiratory tract and establishment of the disease [1–3].

ACT belongs to an extensive family of T1SS-secreted toxins of Gram-negative pathogens, referred to as RTX (Repeats in ToXin) family, characterized by the presence in the C-terminal end of their sequences of numerous calcium-binding sites formed by Gly- and Asp-rich nonapeptide repeats [4,5]. ACT is a 1706 amino acid polypeptide initially synthesized as pro-toxin, that is covalently acylated in the bacterial cytosol at two conserved internal Lys residues (Lys 860 and Lys 983) by a dedicated acyltransferase, CyaC [6]. ACT is then secreted across both bacterial membranes by the type I secretion system [7]. Calcium binding to the RTX repeats (at mM range) and post-translational palmitoylation promote folding of ACT making the toxin fully competent for biological activity [8].

ACT is distinguished from the rest of RTX toxins by bearing a cell-invasive N-terminal enzymatic adenylate cyclase (AC) domain (~364 residues) fused to a C-terminal RTX haemolysin moiety (~1342 carboxy-proximal residues) [5]. The catalytic AC domain converts ATP into cAMP [9]. The C-terminal RTX moiety is responsible for the translocation of the AC domain across the host plasma membrane and for the lytic properties of ACT on cells [5]. This RTX moiety further consists of: a translocation region (TR), spanning residues ≈400 to 500, which has been directly involved in the transport of the AC domain across the plasma membrane [10]; a hydrophobic domain (HD), spanning residues ≈500 to 700, containing several α-helical segments, which have been involved in pore formation [11]; an acylation region spanning residues 750 to 1000 that contains the two conserved acylation sites [6]; a calcium-binding RTX domain, between residues 1008 and 1590, which harbours the characteristic Gly- and Asp-rich nonapeptide repeats that form the numerous (~40) calcium-binding sites of ACT— hallmark of ACT membership to the RTX family [4]. The last ≈116 residues at the RTX haemolysin domain constitute the non-cleavable secretion signal recognized by the dedicated T1SS system [12].

It is believed that the primary ACT targets are myeloid phagocytic cells that possess the CD11b/CD18 integrin, which acts as toxin receptor [13], although ACT can also efficiently intoxicate a variety of cells lacking the integrin, such as erythrocytes or epithelial cells, likely through a direct interaction with their plasma membrane [14]. To generate cAMP inside the target cell, ACT binds to the cell membrane and translocates its AC domain across the plasma membrane by a mechanism that remains poorly understood. Once in the cytosol, the AC domain catalyzes the unregulated conversion of intracellular ATP to cAMP in a reaction that is stimulated by eukaryotic calmodulin [9]. Production of unregulated levels of cAMP subverts cellular physiology and suppresses bactericidal functions of phagocytes [15]. The RTX haemolysin moiety of ACT forms in lipid bilayers oligomeric pores that account for the haemolytic activity of the toxin [16, 17].

The exact step-by-step membrane interactions of ACT leading to AC domain translocation and to formation of lytic pores remains poorly understood, due in part to lack of structural data of membrane-inserted toxin molecules. From different mutational studies it has been proposed that the TR and the HD of ACT are directly implicated in AC domain translocation and pore formation [11, 18–25]. Both TR and HD are predicted to consist of several α-helices. In the TR two α-helices are predicted to form between residues 413 to 434 and 454 to 484, respectively that appear to interact with lipid bilayers [11, 25]. In the HD five putative amphipathic or hydrophobic α-helices (HI_502–522_, HII_527–550_, HIII_571–592_, HIV_607–627_ and HV_678–698_) have been traditionally considered (predicted by the algorithm of Eisenberg) [26].

By analogy with other known pore-forming toxins, it is assumed that the hydrophobic helical domain of ACT inserts into the lipid bilayer forming a hydrophilic pore that would eventually lead to cell lysis. By contrast, the molecular mechanisms and the structural elements involved in AC domain translocation are less clear, and remain a matter of intense research. Several models have been postulated the last years [27, 28]. One model posits that the ACT pore-forming activity is not implicated in the delivery of the AC domain across target cell membrane, and that on its way into the cytosol the translocating AC domain bypasses the cation-selective and lytic pore formed by ACT into the membrane [19, 27]. Further, the authors of the model propose that cell-invasive and pore-forming activities of ACT are independent and mutually excluding, operating in parallel in target cell membrane, and predict that two distinct ACT conformers insert into the membrane in parallel, one being the translocation precursor, accounting for AC delivery across the cellular membrane, the other being a pore precursor eventually forming an oligomeric pore and provoking potassium efflux from cells [19]. So far however, none of those supposed two ACT conformers have been isolated. Furthermore, structural integrity of all the transmembrane helices of the HD were shown to be essential for AC domain translocation across the plasma membrane of both CD11b+ and CD11b-cells [19, 23, 24]. A second model posits that upon ACT insertion into the target cell membrane, a helical peptide extending from residue 454 to 484 interacts with the plasma membrane and destabilizes the lipid bilayer which would favour direct AC translocation across the lipid bilayer [28]. High affinity binding of that segment with calmodulin in the cell cytosol would then assist the irreversible translocation of the entire AC domain [28]. Data by others demonstrating the translocation of unrelated polypeptides fused to the ACT haemolysin moiety [29], weaken however reliability of this hypothesis. Recently we showed that an ACT phospholipase A activity might be involved in AC translocation facilitating insertion of transmembrane segments through toroidal perturbations formed by the enzymatic end-product [30].

Our laboratory has recently provided the first nanoscale pictures of ACT lytic pores in lipid membranes [17]. We have revealed that ACT pores are not fixed-sized narrow pores as was believed, but instead are dynamic proteolipidic pores (toroidal pores) involving lipids, besides toxin molecules. Additionally, we have found that cholesterol in the membrane notably stimulates the lytic activity of ACT [31]. These data strongly suggested that ACT might directly interact with this sterol in the membrane. Because several proteins known to directly interact with cholesterol possess in their transmembrane domains sequences the so-called cholesterol-recognition motifs (CRAC motifs with the **L/V**-X(1–5)-**Y/F**-X(1–5)-**R/K** pattern, or CARC motifs with the **R/K-**X(1– 5)-**Y/F**-X(1–5)-**L/V** pattern [32–34] we decided to take a closer look at possible functional cholesterol-recognition motifs in ACT. Here we have identified 38 CRAC and CARC putative motifs. Basing on their specific location in ACT sequence we have focused our investigation on four of such motifs, namely: the CARC^415^, CRAC^485^, CRAC^521^ and CARC^532^ sites located between amino acids 400-700. We reveal that the four motifs are real, functional cholesterol-binding sites, and crucial for both lytic and translocation activities of ACT on cells.

## RESULTS

### Numerous potential cholesterol-recognition motifs can be identified in ACT sequence

Firstly we searched the ACT sequence for CRAC and CARC motifs. The sequence of ACT was obtained in FASTA format from UniProt (http://www.uniprot.org/). A search for CRAC and CARC motifs was then performed with EMBROSS: fuzzpro program (http://emboss.bioinformatics.nl/cgi-bin/emboss/fuzzpro). Sequences given as a search pattern were: [LV]-X(1,5)-**Y/F**-X(1,5)-[RK], [RK]-X(1,5)-**Y/F**-X(1,5)-[LV]. We identified 20 possible CRAC motifs and 18 potential CARC motifs in the ACT sequence (**Table I**).

**Table I.**
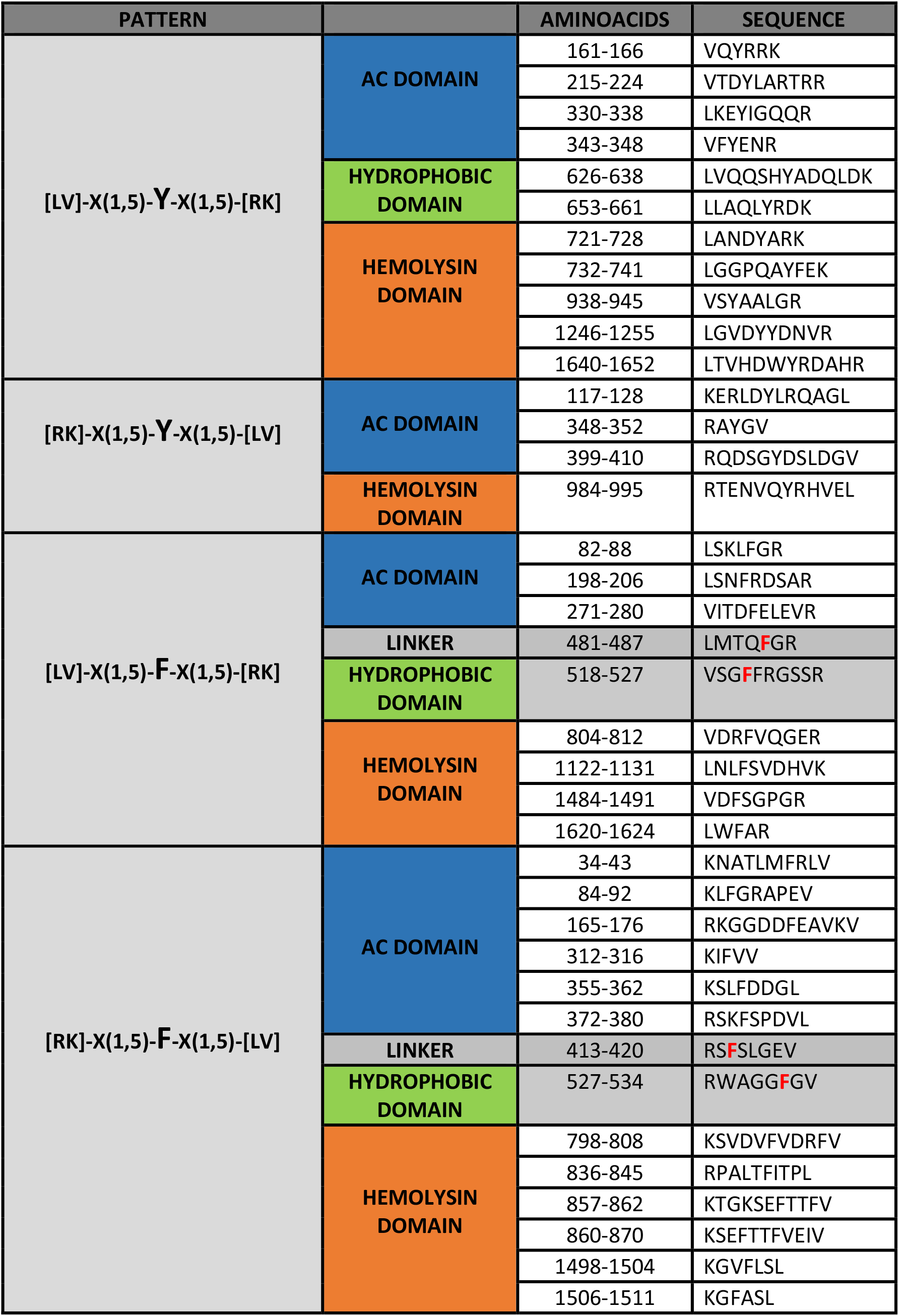

From this set of 38 potential sites we chose four, designated from now on as CARC^415^, CRAC^485^, CRAC^521^ and CARC^532^, for further investigation. This selection was based on the particular location of these four sites within ACT (**Fig 1**), the CARC^415^ and CRAC^485^ motifs are localized in the TR (residues ≈400-500) and the CRAC^521^ and CARC^532^ sites are in the HD (residues ≈500-700), both supposed to interact and insert into the target cell membrane and to be important for ACT functionality. The CARC^415^ motif (residues 413-420, RS**F**^415^SLGEV) is located at the N-terminus of a long α-helix (h1, from now on) predicted to form between residues ≈413 to 434 of the TR, while the CRAC^485^ motif (residues 481-487, LMTQ**F**^485^GR) is located at the C-terminus of a second long α-helix (h2, from now on) predicted to form between residues ≈454 to 484 of this region [27]; On other side, the CRAC^521^ motif (residues 518-527, VSG**F**^521^FR) localizes at the C-terminus of the first predicted α-helix (HI, from now on) of the HD, while the CARC^532^ motif (residues 527-534, RWAGG**F**^532^GV) is located at the N-terminus of the second α-helix (HII, from now on) of the HD. Thus, we anticipated that the selected four potential CRAC and CARC motifs might be relevant mediating the toxin interaction with the cell membrane cholesterol, and decided to test them.

**Figure 1.**
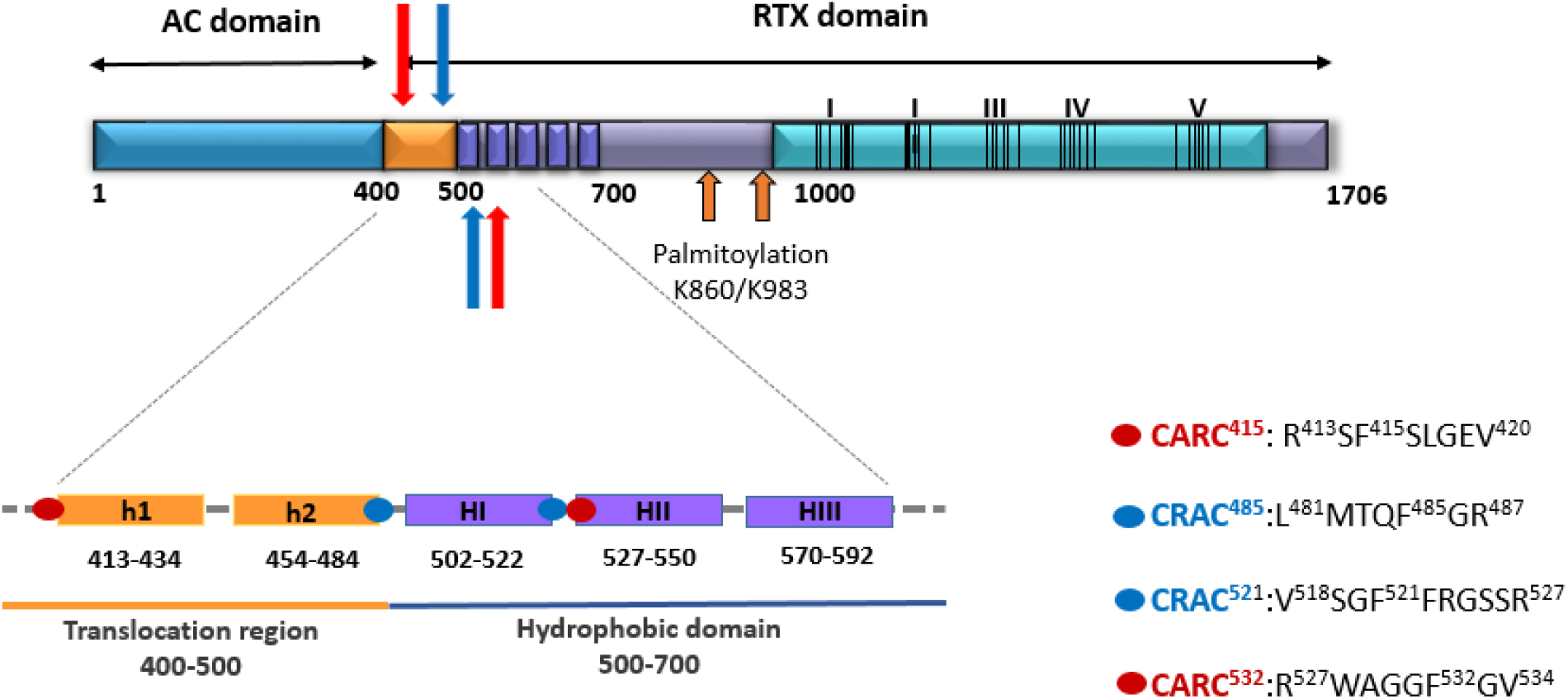
Schematic drawing of the ACT polypeptide chain in which the investigated four potential cholesterol-recognition motifs are detailed. Two predicted α-helices in the TR, namely, h1 and h2, and three of the five predicted amphipathic and hydrophobic helices of the pore-forming domain, namely HI, HII and HIII have been depicted with more detail. Blue or red spots have been used to specify the location of each one of the four potential cholesterol-recognition motifs in each of the helical segments.

### Preincubation of ACT with free cholesterol, or extraction of the sterol from the cell membrane with methyl-β-cyclodextrin inhibit the toxin-induced haemolysis

To corroborate that cholesterol plays a role in membrane binding and haemolysis by ACT, the toxin (100 nM) was preincubated with increasing concentrations of free cholesterol for 30 minutes at RT, and was then further incubated with erythrocytes in the presence of free cholesterol, and haemolysis was measured.

As depicted in **Fig 2A**, free cholesterol at concentrations above 5 μM had a notable inhibitory effect on the toxin-induced erythrocyte lysis, suggesting that ACT may recognize and directly bind to membrane cholesterol. This idea was reinforced by other experiment in which cholesterol was extracted by pretreatment for 30 min of erythrocytes with methyl-β-cyclodextrin (5 mM), an agent commonly used to remove cellular cholesterol [35]. As illustrated in **Fig 2B**, depletion of cholesterol by this compound reduced importantly the toxin-induced haemolysis in an ample range of toxin concentrations. Together these results suggested thus that ACT might have one or more specific binding sites for the sterol.

**Figure 2.**
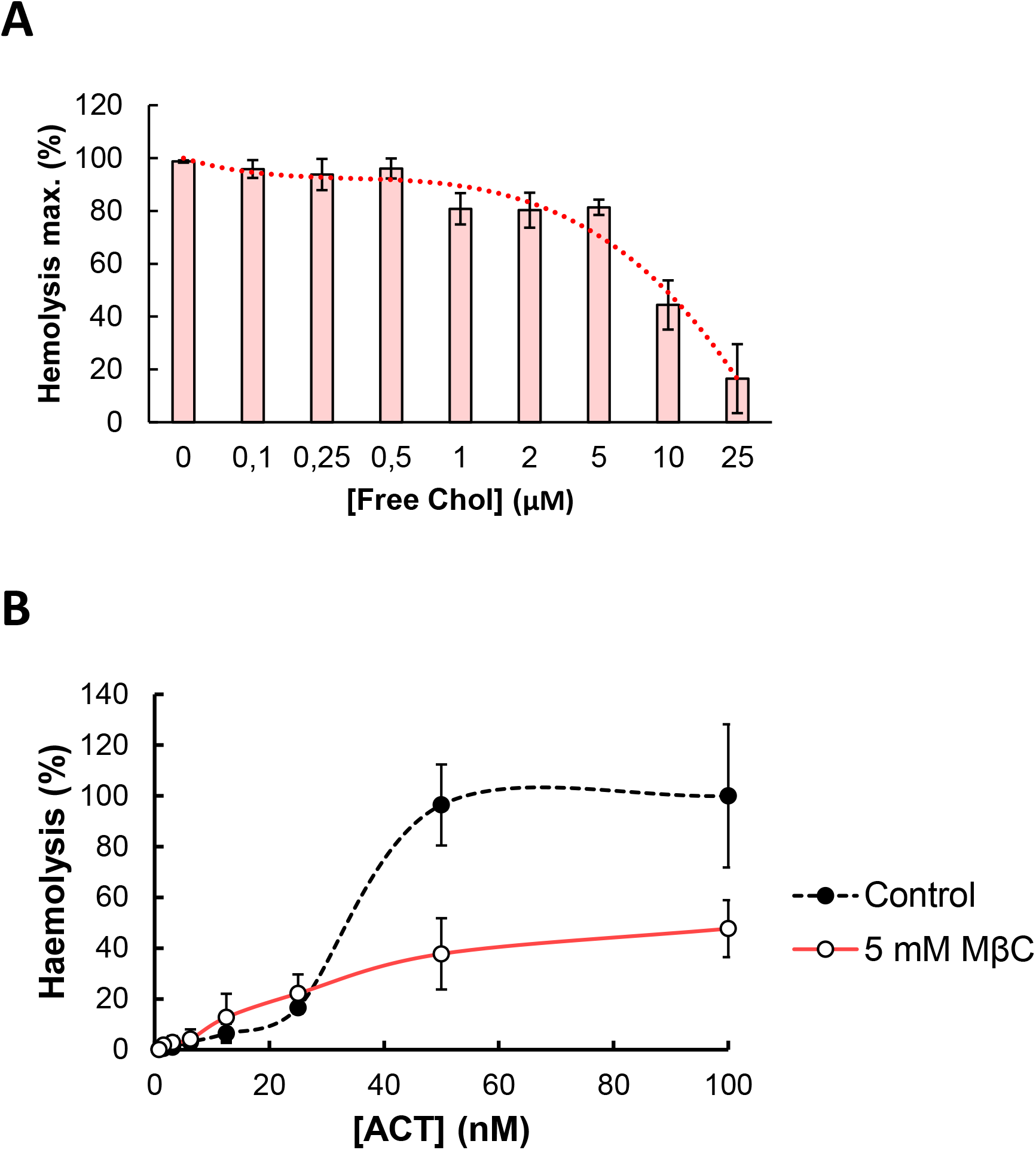
Effect of ACT preincubation with free cholesterol and of membrane sterol depletion on ACT-induced haemolytic activity. (**A**) ACT (100 nM) was preincubated for 30 minutes at RT in the presence of free cholesterol (0-25 μM). Then sheep erythrocytes at a density of 5 × 10^8^ cells/ml were added and the mixture was further incubated for 180 min at 37°C. Haemolytic activity was measured as decrease of turbidity at 700 nm and expressed as haemolytic percentages (calculated as detailed in the Experimental Procedures section). Data represented in the figure correspond to the mean of three independent experiments ±SE. (**B**) Sheep erythrocytes (5 × 10^8^ cells/ml) were pre-treated with methyl-β-cyclodextrin (5 mM) for 30 min at 37°C to decrease the cholesterol content available in the cell membrane. Then ACT was added at different concentrations and further incubations of 180 min were performed before determining the lysis percentage.

### Substitutions by Ala of the central Phe in 415, 485, 521 and 532 positions, in the respective potential cholesterol-recognition motifs of ACT, have a differentiated effect on the toxin-induced haemolysis

Mutations in the central Tyr or Phe residues in CRAC and CARC motifs have been shown to strikingly reduce or eliminate protein-cholesterol interactions in different cholesterol-binding proteins, affecting consequently protein activity in membranes [32–34].

To determine whether the here selected four sites are indeed functional, and to examine their possible implication in cholesterol binding by ACT, we constructed several mutant proteins with single Ala substitutions in the central Phe residues 415, 485, 521 and 532 of the respective motifs. Then we checked firstly the effect of these mutations on the toxin-induced haemolysis.

**Fig 3** shows the raw traces of the kinetics recorded from a representative experiment of haemolysis induced by wild-type ACT (50 nM) or by each one of the four mutant toxins (50 nM), namely F415A, F485A, F521A and F532A mutants. From haemolytic kinetics such as the observed in **Fig 3**, maximum haemolysis percentages were obtained at 180 min and were represented in **Fig 4A**. In addition, the t 1/2 values (time required to induce 50% haemolysis) were plotted in **Fig 4B**.

**Figure 3.**
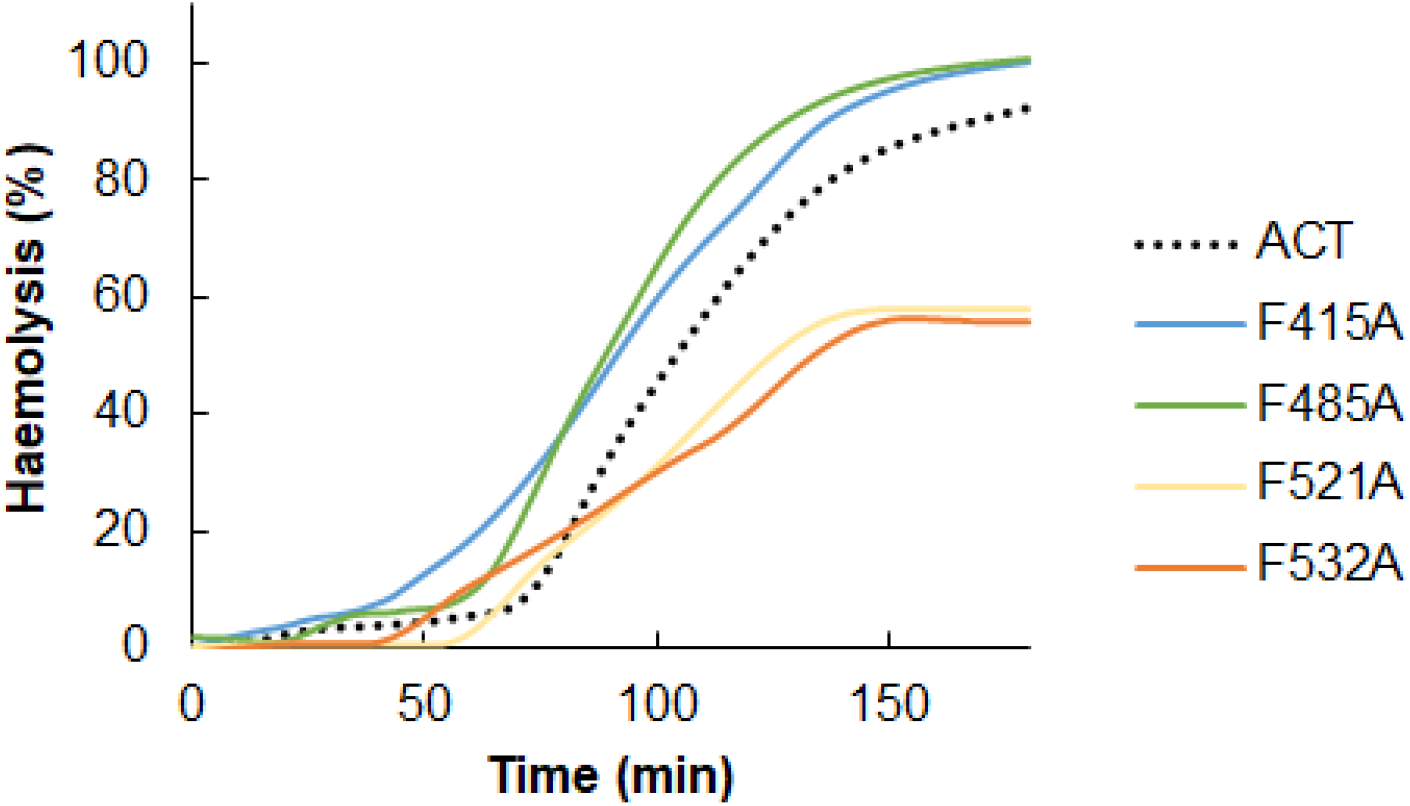
Effect of point Ala substitutions in the central Phe residues of the potential cholesterol-binding sites CRAC^415^, CARC^485^, CRAC^521^ and CARC^532^ on the kinetics of the ACT-induced haemolysis. Raw traces of the kinetics recorded from a representative experiment of the haemolysis induced by intact ACT (50 nM) or by each one of the four mutant toxins (50 nM). A suspension of sheep erythrocytes (5 × 10^8^ cells/ml) was incubated with each protein for 180 min at 37°C, recording the scattering changes measured at 700 nm at every second. Then the haemolysis percentage was calculated as detailed in the Experimental Procedures section and depicted in the figure. The traces shown correspond to a representative experiment from three experiments performed independently.

**Figure. 4.**
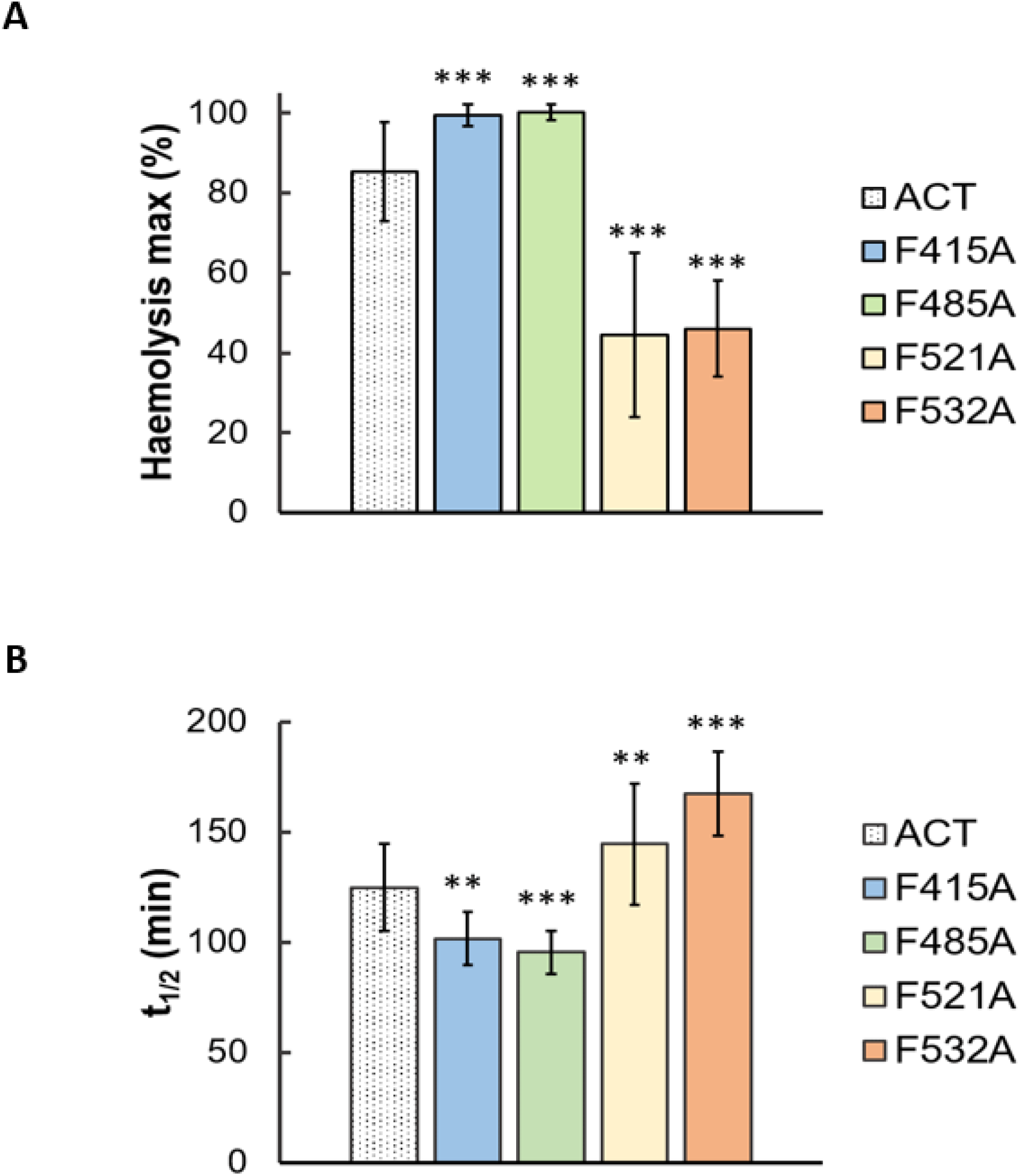
Effect of point Ala substitutions in the central Phe residues of the potential cholesterol-binding sites CRAC^415^, CARC^485^, CRAC^521^ and CARC^532^ on the (A) maximum haemolytic percentage, and (B) t_1/2_ of the ACT-induced haemolysis. Haemolysis induced by 50 nM of intact ACT or by each one of the four mutant toxins was assayed with a suspension of sheep erythrocytes (5 × 10^8^ cells/ml) incubated with each protein for 180 min at 37°C. Data represented in the figure correspond to the mean of three independent experiments ± SE.

As observed in the figures, the effect of the mutations was different depending on the location of the CRAC/CARC sites in the ACT primary structure. The individual substitutions of Phe by Ala in the respective CRAC^521^ and CARC^532^ motifs (F521A and F532A) at the HD, induced a prominent inhibitory effect in the lytic activity of the mutant toxins, in both cases slowing down the erythrocytes lysis (**Fig 3 and Fig 4B**) and reducing to the half the maximum haemolysis extent after 180 min of incubation (**Fig 4A**). In contrast, the single Ala substitutions of the central Phe in the CARC^415^ and CRAC^485^ motifs (F415A and F485A), led to a faster and a greater lytic activity of the respective mutant toxins, reflected in the significantly greater maximum haemolysis values obtained after 180 min incubation, and in the lower t 1/2 values (time in minutes required to induce 50% haemolysis) (**Fig 4**)

To check whether such mutations had any effect on toxin binding to lipid bilayers we performed a control experiment. Data represented in **Fig 5** indicated that the binding percentage was similar for the four mutant toxins relative to the intact ACT. This allowed us to rule out that the inhibition in the lytic activity caused by the F521A and F532A mutations was due to a lower protein binding. Similarly we could discard a greater binding as possible cause of the observed increment in the haemolysis percentage observed for the F415A, F485A mutant toxins. Together these data were a strong indication that the four cholesterol sites explored were real, functional sites, and that, while sterol binding through the CRAC^521^ and CARC^532^ motifs sites is essential for the pore-forming activity of ACT, the interaction of CARC^415^ and CRAC^485^ motifs with membrane cholesterol seems to hinder the ACT lytic activity, since preventing that interaction with the sterol by the F415A and F485A mutations promotes a greater lytic activity.

**Figure 5.**
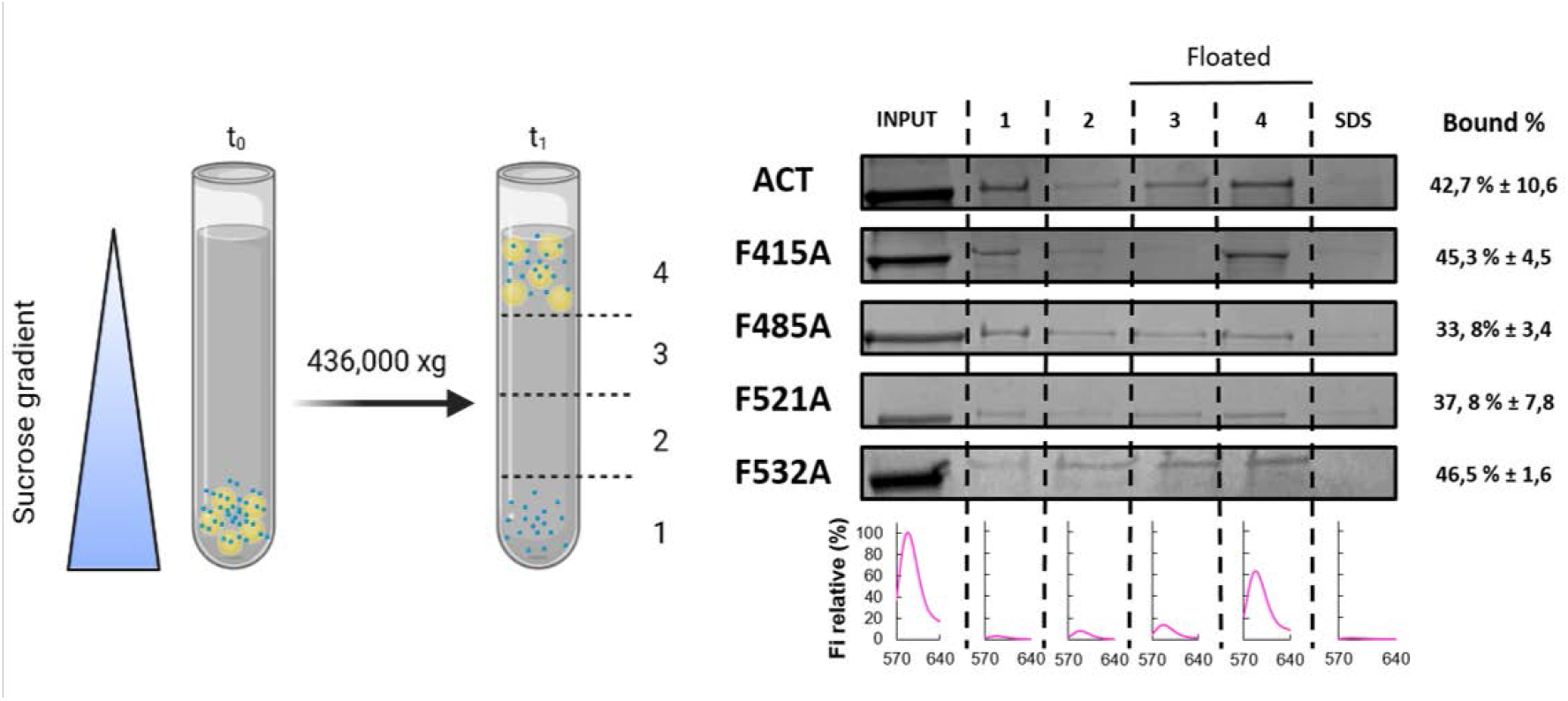
Quantification of the binding of ACT or ACT mutants to lipid bilayers. Membrane partitioning as measured by flotation assays using large unilamellar vesicles composed of DOPC:Chol (3:1 molar ratio). Details on the flotation assay methodology are provided in the section of Experimental Procedures.

### Substitutions by Ala of the central Phe in 415, 485, 521 and 532 positions, in the respective cholesterol-recognition motifs of ACT, inhibit prominently AC domain translocation

To determine whether the CARC^415^, CRAC^485^, CRAC^521^ and CARC^532^ motifs have any role in AC domain delivery, we measured the effect on cAMP production of the Ala substitution in the central Phe residues 415, 485, 521 and 532 of the respective mutant proteins, in J774A.1 cells (**Fig 6**).

**Figure 6.**
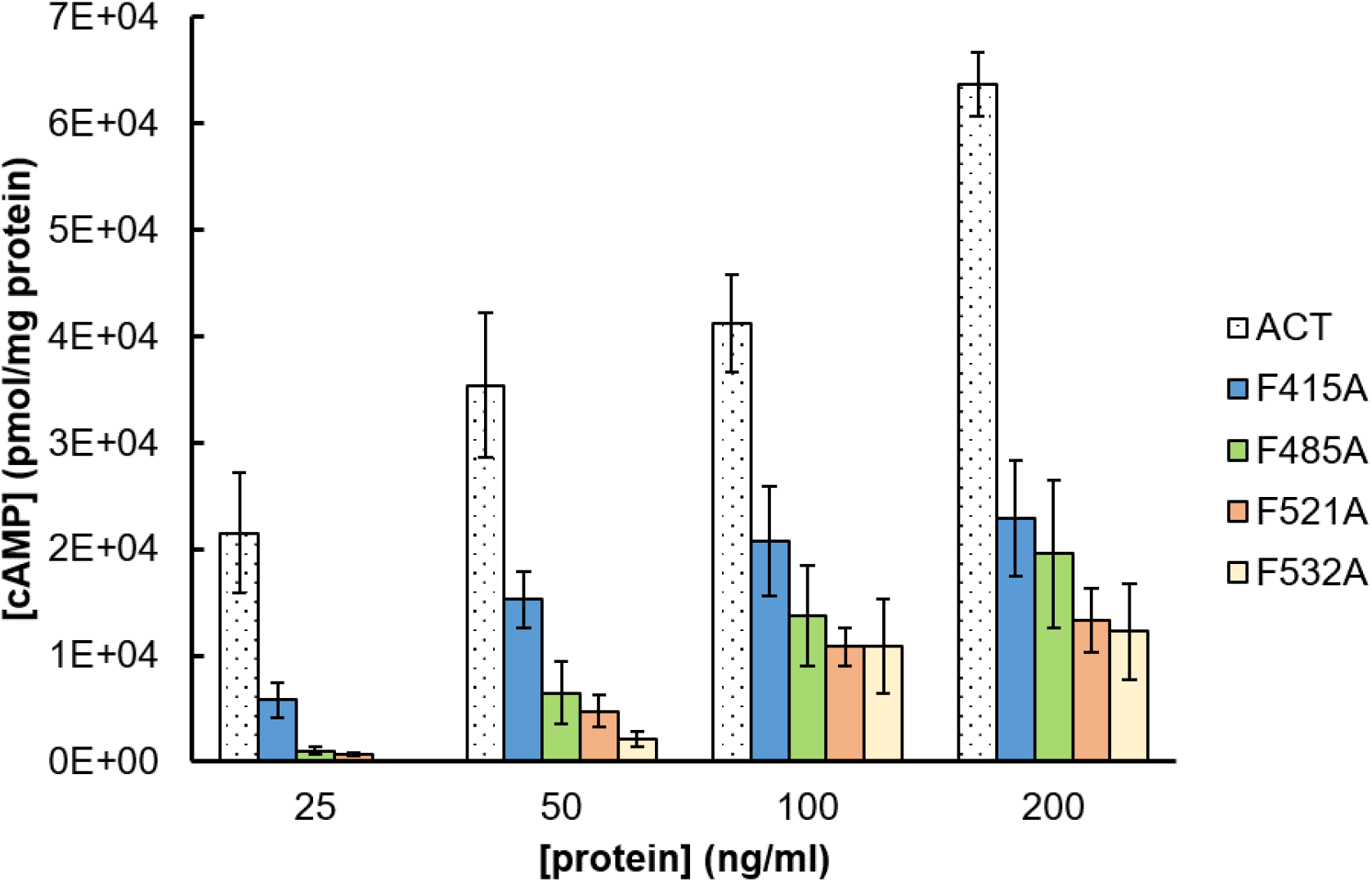
Point Ala substitutions in the central Phe residue of the potential cholesterol-binding sites CRAC^415^, CARC^485^, CRAC^521^ and CARC^532^ prominently decrease AC domain translocation. Translocation of AC domain was assessed by determining the intracellular concentration of cAMP (pmol/mg protein) generated in J774A.1 cells (1x 10^5^ cells/ml) suspended in 20 mM Tris-HCl, pH=8.0 buffer supplemented with 150 mM NaCl and 2 mM CaCl_2_, upon treatment for 30 minutes at 37°C with different concentrations (25-200 ng/ml) of intact ACT or the corresponding mutant toxin.

As reflected in **Fig 6**, relative to intact ACT, the four single mutations F415A, F485A, F521A and F532A greatly impacted the capacity to deliver the AC domain by the respective mutant proteins in a range of toxin concentration between 25-200 ng/ml, with a more prominent inhibitory effect observed for the F521A and F532A mutations. Given that none of the mutations had any significant effect on toxin binding to cholesterol-containing lipid bilayers, results in **Fig 5** corroborated the previous conclusion that the selected four potential cholesterol binding motifs are real and functional cholesterol binding sites, and indicated besides, that ACT interaction with membrane cholesterol through the CARC^415^, CRAC^485^, CRAC^521^ and CARC^532^ motifs is crucial for AC domain translocation.

## DISCUSSION

In this study we have made two major observations. First, we show that ACT binds free cholesterol, and identify in ACT sequence 38 potential cholesterol-recognition motifs, distributed all along the toxin primary structure. Second, we reveal that four of those motifs are real, functional cholesterol-binding sites, since substitutions by Ala of their central Phe residues have important consequence on the lytic and translocation activities of ACT on target cells, suggesting their direct intervention in cholesterol recognition and toxin functionality.

We find that single mutation by Ala of the central Phe in 521 and 532 residues in the CRAC^521^ and CARC^532^ motifs, respectively, causes a potent reduction in the lytic and translocation capacities of ACT, without affecting its membrane association. Given that those motifs are placed in the HI and HII helices of the pore-forming domain (**Fig 1**), it can be inferred that a direct, and likely specific, interaction between those motifs and cholesterol drives the membrane insertion of these two helices, a step that, as judged by the results, is essential, for both haemolysis and AC translocation. We find, as well, that in the F415A and F485A mutants the AC translocation capacity is notably inhibited, whereas simultaneously they exhibit a greater lytic activity. Since the CARC^415^ and CRAC^485^ motifs are in the h1 and h2 helices of the TR, a region reported to interact with the membrane and to modulate the ACT pore-forming activity [11, 25], we can infer that cholesterol binding through these two motifs favours the insertion of h1 and h2 into the lipid bilayer. Our data indicate also that, such interaction, being necessary for AC domain translocation, simultaneously hinders the ACT lytic activity. Together thus, our results not only reaffirm the pivotal role played by the h1, h2, HI and HII helices in ACT biological activity, but, additionally, as we will crumble in the next paragraphs, they allow to delineate a plausible membrane topology for each helix in the ACT activities.

The CARC^532^ motif (R^527^WAGGF^532^GV^534^) does not contain any polar charged residue except the N-terminal R527, and is located at the N-terminus of the HII helix (residues 529-549), which is one of the hydrophobic α-helices in the ACT pore-forming domain predicted to be transmembrane [22–24]. It is thus very likely that HII helix adopts a transmembrane position and that the N-terminal CARC^532^ motif is embedded into the membrane. To predict its most likely orientation into the bilayer (N→ C or C→ N), we will assume the prevalence of the empirically demonstrated “positive-inside rule” (preferential occurrence of positively charged residues (Lys and Arg) at the cytoplasmic edge of transmembrane helices) [36, 37] in transmembrane helices and that both CARC and CRAC motifs are lineal oriented motifs. Then, it can be envisaged that HII helix would insert with its N-terminus oriented to the cytosolic side and the C-terminal end to the extracellular side of the membrane (C→ N orientation). This orientation would place the R^527^ residue and its positively charged guanidinium group emerging on the cytosolic side of the membrane (snorkelling effect) [38], while the hydrophobic V^534^ would remain buried into the bilayer.

Upstream and adjacent to HII is the amphipathic HI helix (residues ≈502-522) and the CRAC^521^ site (V^518^SGF^521^FR^523^) at its C-terminus. HI possesses two Glu residues (E^509^ and E^516^) in the middle of its sequence, so its transmembrane insertion would be poorly favourable. However, it is expected that a thermodynamically favourable binding to membrane cholesterol through the CRAC^521^ motif could drive HI insertion into the membrane. Given the C→ N orientation of HII, then HI would insert with N→ C orientation. This would place the two CRAC^521^ and CARC^532^ motifs to the cytosolic side of the membrane, and the positively charged cationic groups of the R^523^ and R^527^ side chains emerging at the surface of the inner leaflet. Importantly, HI and HII are separated by the GSS triad (residues 524 to 526), so it is conceivable that HI-HII form a helical hairpin whose transmembrane topology will be greatly stabilized by the cholesterol binding through the CRAC^521^ and CARC^532^ motifs. Due to their sequential proximity, insertion of the HI-HII hairpin will reasonably determine the insertion of the neighbour HIII, HIV and HV helices at the pore-forming domain. Proper transmembrane insertion of all these helices will be implicitly necessary to form a functional pore structure. It is thus easily envisaged that mutations that affect cholesterol binding in either of the mentioned CRAC or CARC motifs, would have a deleterious, destabilizing effect on the hairpin insertion, which can explain pretty well the potent inhibitory effect of the F521A and F532A mutations, on the ACT lytic capacity as shown here. On other side, given the C→ N orientation of HII helix, then the downstream HIII helix (residues ≈570-594) would insert with its N-terminus oriented to the extracellular side and the C-terminus towards the cytoplasmic side, positioning most likely the negatively-charged E^570^ at the extracellular side, while the R^594^ would locate at the cytoplasmic side of the membrane (positive-inside rule) [36].

Preceding the HI helix is the TR, constituted by the h1 and h2 helices, each one of which has a cholesterol-recognition motif in one end. It can thus be envisioned that a favourable cholesterol binding through the CRAC^415^ and CARC^485^ motifs could drive the h1 and h2 insertion into the membrane. Because the CRAC^415^ site (R^413^SF^415^SL^417^GEV^420^) is at the N-terminus of h1, and the CARC^485^ site (L^481^MTQF^485^GR^487^) at the C-terminus of h2, and the downstream HI inserts with a N→ C orientation, then it can be anticipated that cholesterol binding by these two sites would favour h2 insertion with its C-terminus to the extracellular side and the N-terminus at the cytosolic side, and h1 insertion with its N-terminus to the extracellular side and the C-terminus to the cytosolic side. The h2 helix was predicted by other groups to be a long α-helix extending from residues ≈454 to 484, placing the two positively-charged Arg residues, R^461^ and R^474^, in the middle of the helix [25, 39], which would presumably make very unfavourable its transmembrane insertion. However, the presence of the C-terminally located CARC^485^ motif and its binding to cholesterol could expectedly turn the transmembrane insertion of this h2 thermodynamically favourable. This transmembrane topology of h2 would be further favoured by placing the positively charged R^461^ at the cytosolic side of the membrane, and perhaps the R^487^ (last residue of the CARC^485^ motif) at the extracellular side. This would make the h2 helix a little bit shorter at the N-terminus and a little bit longer at the C-terminus (residues 461 to 487) as compared to the length predicted for this helix by other investigators [25, 39], and would locate a single positive residue, the R^474^, within h2. In the case of h1, it is expectable that it would be flanked by the positively charged R^435^ residue at the cytosolic side, and the R^413^ at the extracellular flank. In sum, binding to membrane cholesterol emerges as instrumental for the proper membrane topology of both, the TR and the HD, and consequently essential for the toxin functionality. A scheme of the complete membrane topology for the h1, h2, HI, HII and HIII helices, as predicted here, has been drawn in **Fig 7**.

**Figure 7.**
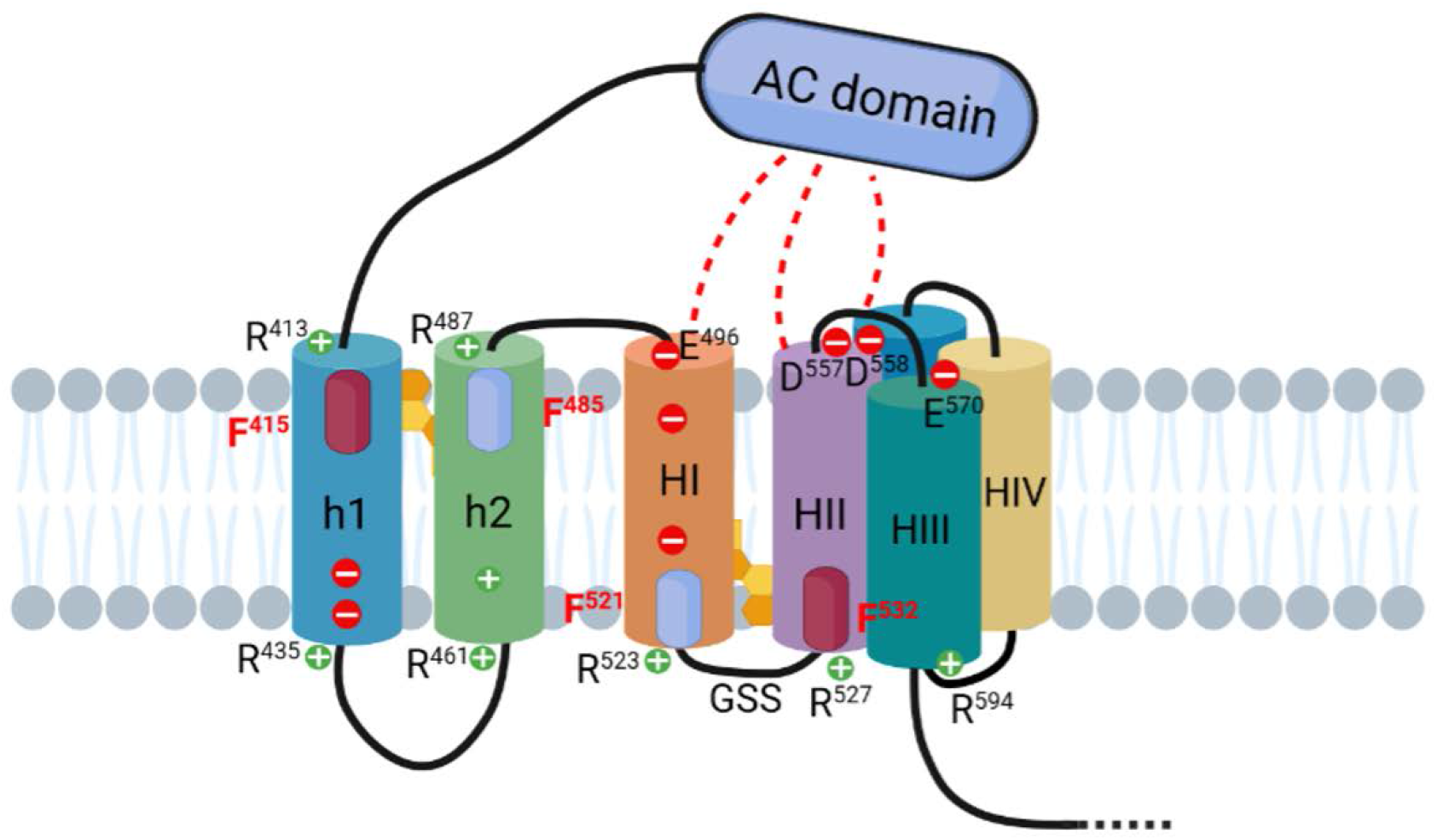
Schematic model of the proposed membrane topology for the helical elements constituting the TR and the HD of ACT. The figure shows a scheme of the complete membrane topology for the h1, h2, HI, HII and HIII helices, as predicted here, on the basis of the experimental data shown in our study. From the proposed topology it can be concluded that both for AC transport and for lytic activity, ACT would adopt a single transmembrane topology as drawn here, that is stabilized by the binding to membrane cholesterol through the here identified four CRAC/CARC motifs. We hypothesize that the cholesterol-mediated transmembrane insertion of the h1-h2-HI-HII helices would bring the extracellularly located AC domain near the pore structure. Spatial proximity of the AC domain from the pore would plausibly allow interactions to be stablished between segments of the AC domain and one or several residues of said helices forming the pore. Such native interactions would be necessary to assure penetration of the AC polypeptide into the pore lumen and its transport to the target cytosol. The h1 and h2 helices localize to the so-called TR of ACT extending from residues ≈400-500. HI to HV helices form part of the HD which extends from residues ≈500-700. The four CARC and CRAC motifs identified, described in the text, have been drawn in the model as magenta and blue colour ellipses, respectively. Red dashed lines between the AC domain and helices of the HD represent possible native interactions that would expectedly be stablished and that would be necessary to spatially approach the AC domain to the entrance of the pore formed by the hydrophobic helices. More details are given in the text of the **Discussion** section.

Of note, ACT contains at its HD other four CRAC motifs (CRAC^632^, CRAC^658^, CRAC^725^ and CRAC^738^), all of which have a central Tyr residue instead of Phe (**Table I**). However, as recently reported, none of them appears to be involved in cholesterol recognition by ACT [40], which sounds consistent with their location in extracellular segments between helices of the HD. Therefore, by warranting the proper intra-membrane topology of the h1, h2, HI-IV helices, the here identified four cholesterol-recognition motifs would represent a “cholesterol sensor” necessary to initiate membrane insertion of two ACT regions essential for the toxin biological activities. Existence of such molecular mechanism may explain pretty well the cholesterol dependency shown by these ACT activities on target cells [32].

In the absence of structural data, the transmembrane topology and organization of the ACT translocation and HD involved in both AC delivery and pore formation has remain elusive. For more than two decades it has been assumed in the field that the pore-forming and the AC translocating activities associated with the ACT C-terminal haemolysin moiety (residues ≈400-1706) are fully independent, and occur in parallel, being associated with two different toxin conformers, one that would lead to direct AC transport across the lipid bilayer, and other, that upon oligomerization, would lead to pore formation [18–20, 40]. However, so far no demonstration has been provided for the existence of these hypothetical two different ACT conformers. Instead, on the basis of the here shown experimental data, and the membrane topology delineated from them, it can be concluded that both for AC transport and for lytic activity, ACT would adopt a single transmembrane topology, stabilized by the binding to membrane cholesterol through the here identified four CRAC/CARC motifs. These data challenge thus the previously accepted model of conformational duality of ACT to perform its two biological activities [19, 40].

For long it has been also believed that the ACT pores are too small (0.6–0.8 nm in diameter) for the passage of even an unfolded polypeptide chain, which directly led to discard the possibility that the pore formed by this toxin might serve to transport the AC domain to the target cytosol, and to propose a unique “direct” transport of the AC domain across the plasma membrane [11, 31, 32]. Contrasting with this view, more recent results from our own laboratory have revealed that the ACT pores are of proteolipidic nature, involving lipid molecules besides segments of the protein [17]. As consequence of this more dynamic structure of the ACT pore, it may thus be envision that its hydrophilic lumen can be wider than previously believed. Consistently with this, and given that the AC domain is not itself capable of directly interacting with lipid bilayers [10], and AC delivery requires structural integrity of the pore-forming domain, our present results lead to contend that the simplest, most logic and most plausible mechanism by which the 400-residue-long AC polypeptide is transported to the target cell cytosol, is through the hydrophilic “hole” formed by the ACT pore-forming domain.

We hypothesize that the cholesterol-mediated transmembrane insertion of the h1-h2-HI-HII helices would bring the extracellularly located AC domain near the pore structure. Spatial proximity of the AC domain from the pore would plausibly allow interactions to be stablished between segments of the AC domain and one or several residues of said helices forming the pore (**Fig 7**). Such native interactions would be necessary to assure penetration of the AC polypeptide into the pore lumen and its transport to the target cytosol. It is thus anticipated that mutations that affect the cholesterol binding and hence the insertion of the helices, or that hinder the molecular interactions between the AC segments and pore segments will have a direct effect on AC translocation, in full consonance with our present results. That same reasoning predicts as well that the mutations that would inhibit AC translocation, could simultaneously lead to a lytic activity gain, since the same native interactions could sterically hinder the free ion flux through the pore lumen, perhaps until the translocation has finished and the AC domain is cleaved by target calpain, as recently reported by our laboratory [41]. This would explain why apparently ACT is weakly haemolytic relative to other RTX pore-forming toxins such as *Escherichia coli* α-haemolysin [11]. Consistently, we find here that the F415A and F485A mutations inhibit translocation and enhance the lytic activity. Hampering cholesterol binding through the CARC^415^ and CRAC^485^ motifs would provoke a change in the topology of the h1-h2 helices, to be placed extracellularly and would gain in mobility. This would hinder establishment of the native interactions between the AC segments and the pore segments, moving away the AC domain from the pore entrance. This distancing of the AC domain would eliminate the steric hindrance at the pore entrance allowing a free ion flux, which would be detected as an increased lytic activity, at the same time that the AC delivery would result diminished (**Fig 8**).

**Figure 8.**
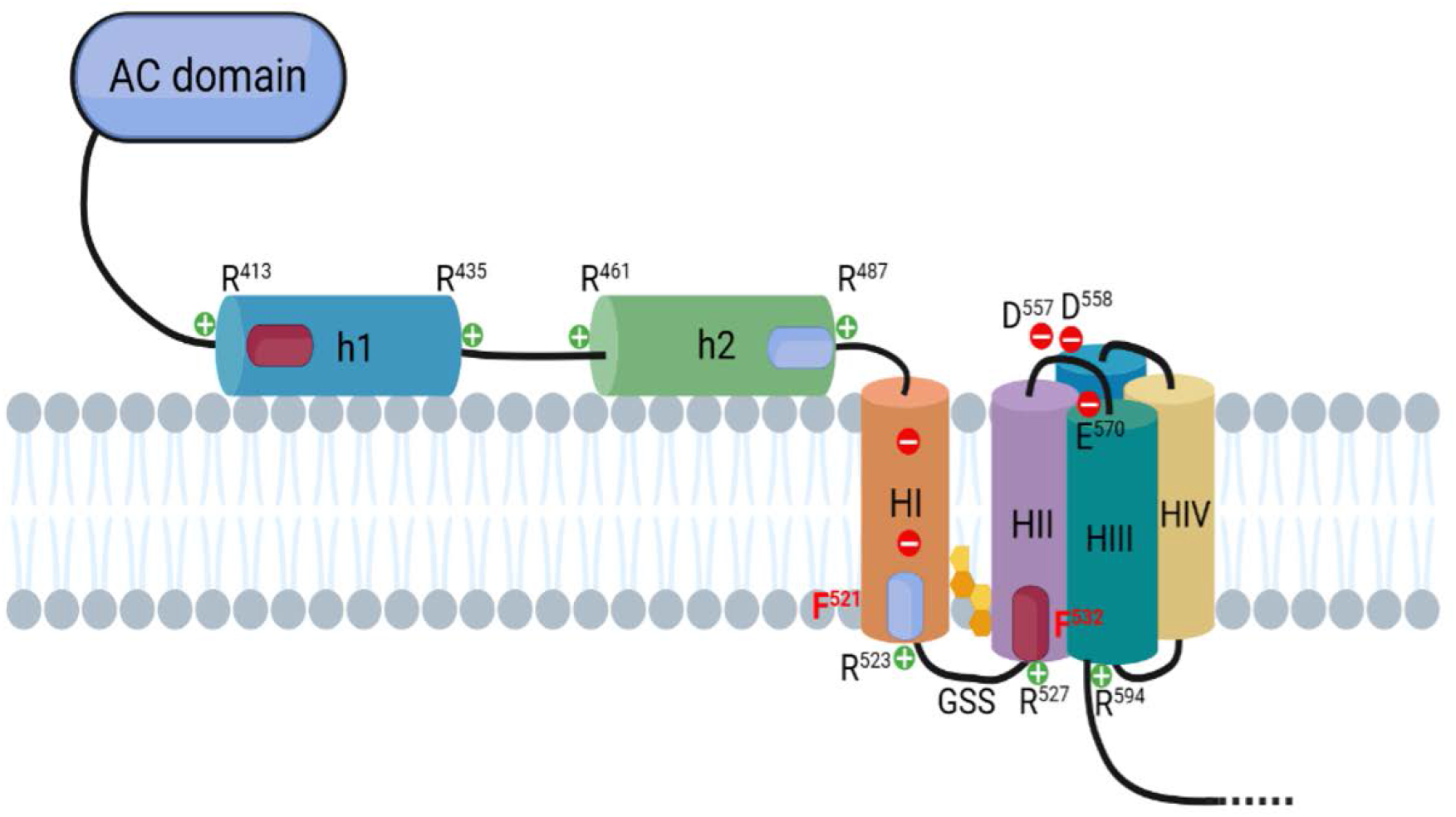
Schematic model of the membrane topology proposed for the helical elements constituting the TR and the HD of ACT for ACT variants in which the Phe in 415 or 485 residues are mutated by Ala. We find here that the F415A and F485A mutations inhibit translocation and enhance the lytic activity. Hampering cholesterol binding through the CARC^415^ and CRAC^485^ motifs would provoke a change in the topology of the h1-h2 helices, to be placed extracellularly and would gain in mobility. This would hinder establishment of the native interactions between the AC segments and the pore segments, moving away the AC domain from the pore entrance, This distancing of the AC domain would eliminate the steric hindrance at the pore entrance allowing a free ion flux, which would be detected as an increased lytic activity, at the same time that the AC delivery would result diminished. More details are given in the text of the **Discussion** section. The four CARC and CRAC motifs identified, described in the text, have been drawn in the model as magenta and blue colour ellipses, respectively.

From our model it is also evidenced the crucial role of the cholesterol binding through the CRAC^521^ and CARC^532^ motifs in ACT activities, since it would allow the intramembrane stabilization of the HI-HII hairpin, which would in turn determine the proper membrane topology of the remaining hydrophobic helices of the pore-forming domain. The HI helix has two negatively charged Glu residues, E^509^ and E^516^, in middle of the helix, which would make transmembrane topology of HI poorly favourable. Cholesterol binding would become thus a way to overcome this energetic penalization, making HI insertion thermodynamically favourable. Curiously, other group had observed that net charge mutations E509K or E516K reduced to the half the AC translocation and cell association, but increased to twofold the haemolytic activity [19]. In contrast, E509V and E509Q substitutions had little effect on toxin activities [19]. Intriguingly, the double substitution E509K+E516K exerted a strong synergic effect. Although the cell association remained similar to that of the single mutants (low binding), the cell-invasive activity of the double mutant was completely abolished, and the haemolytic activity was further enhanced fourfold [19]. Our model predicts that the net charge change in the double mutant (−2 to +2) would inhibit the transmembrane topology of HI, forcing this helix to place out of the membrane. Interestingly, this HI location could change the side of the membrane in which cholesterol would now be recognized by the CRAC^521^ motif, passing from being bound in the cytosolic side of the membrane, to bind it in the extracellular side. Concomitantly, HI exit would force HII to insert into the bilayer with reverse orientation, and to achieve stabilizing through cholesterol binding via the CARC^532^ motif, but now at the extracellular side of the membrane. This topology change of HII would in turn provoke subsequent change in the membrane topology of HIII, and so on for the rest of the helices that conform the pore structure (**Fig 9**). And yet another consequence can be anticipated, the change in location of the flanking residues of each one of the mentioned helices conforming the pore. This way, the residues initially located to the cytosolic side, such as R^527^ (in HII) and R^594^ (in HIII), would move to the extracellular side, whereas the located to the extracellular side, such as S^554^ (in HII), D^557^ and D^558^ (in the loop HII-HIII) and E^570^ (in HIII) would move to the cytosolic side. This would expectedly eliminate the aforementioned native interactions established with segments of the AC domain, moving the AC domain away from the pore entrance and, affecting consequently the AC translocation. Moreover, ion selectivity of the pore could also result altered by the inverted location of the residues at the ends of the helices. Fully supporting this it was detected a drop in the cation selectivity in the pores formed by the E509K+E516K double mutant in black lipid bilayers [19]. On the contrary, the intensification of the haemolytic activity observed in this mutant [19] is somehow perplexing, since it suggests that the ACT pore may be “reversible”, this is, no matter whether it is inserted with a given topology or if it is inserted with the reverse, in both cases the ions seem to be able to flow freely, as long as the entrance is not blocked by the AC domain.

**Figure 9.**
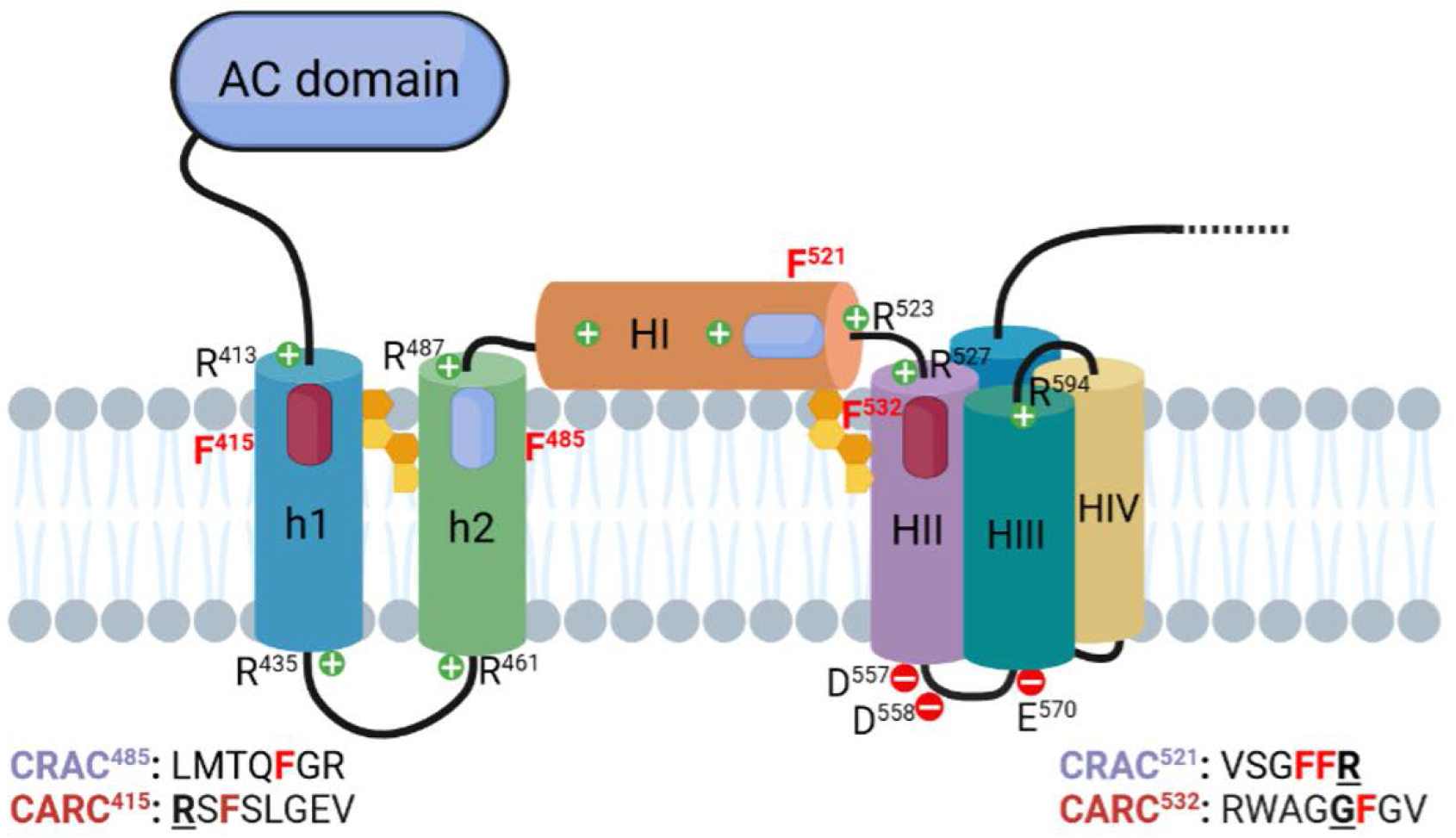
Schematic model of the membrane topology of the helical elements constituting the TR and the HD for the double mutant E509K+E516K. Our model predicts that the net charge change in the double mutant (−2 to +2) would inhibit the transmembrane topology of HI, forcing this helix to place out of the membrane. This HI location could change the side of the membrane in which cholesterol would now be recognized by the CRAC^521^ motif, passing from being bound in the cytosolic side of the membrane, to bind it in the extracellular side. Concomitantly, HI exit would force HII to insert into the bilayer with reverse orientation, and to achieve stabilizing through cholesterol binding via the CARC^532^ motif, but now at the extracellular side of the membrane. This topology change of HII would in turn provoke subsequent change in the membrane topology of HIII, and so on for the rest of the helices that conform the pore structure. This would expectedly eliminate the aforementioned native interactions established with segments of the AC domain, moving the AC domain away from the pore entrance and, affecting consequently the AC translocation. Interestingly, ion selectivity of the pore could also result altered by the inverted location of the residues at the ends of the helices.

In sum, to our best knowledge, the here presented model of membrane topology accounts for all available experimental data and suggests a plausible mechanism by which ACT can translocate the AC domain on target cells, at the cost of sacrificing the lytic potency.

Given the relevance of the specific cholesterol-recognition sites in ACT activity, it can be anticipated that targeting the here identified four CRAC/CARC motifs could be a new therapeutic option for inhibiting cholesterol-binding and hence reducing the toxicity of ACT on cells.

## EXPERIMENTAL PROCEDURES

### Expression and purification of intact ACT

ACT was expressed in *Escherichia coli* XL-1 blue cells (Stratagene) transformed with pT7CACT1 plasmid, kindly provided by Dr. Peter Sebo (Institute of Microbiology of the ASCR, v.v.i., Prague, Czech Republic) and purified as described by Karst et al. [8].

### Construction, expression and purification of the ACT mutants F415A, F485A, F521A and F532A

The variants of ACT F415A, F485A, F521A and F532A were cloned, expressed and purified from *E. coli*. cyaA DNA was amplified from genomic DNA by PCR and cloned in pET-15b (GenScript) using AsuII and NcoI enzymes to generate plasmid pME14. Site-directed mutagenesis according to Agilent protocol was performed on pME14 to replace Ala codons for Phe in 415, 485, 521 and 532 residues. All plasmid inserts were sequenced to confirm accuracy of PCR and mutagenesis. For protein expression, *E coli* BL21 transformed with pME14 plasmid was grown in LB with 100 μg ml^−1^ ampicilin to A_600_ =0.6-0.8 and protein expression was induced by 4h growth in 1 mM isopropyl-β-D-thiogalactopyranoside. Protein purification was performed according to the method described in Karst et al (2014) [8]. Concentrations of purified ACT proteins were determined by the Bradford assay (Bio-Rad, USA) using bovine serum albumin as standard. All toxins purified by this method were more than 90% pure as judged by SDS-PAGE analysis (not shown).

### Haemolysis assay

Haemolysis assays were performed on 96-well plates. Briefly, serial dilutions of ACT (starting at 50 nM) in assay buffer (20 mM Tris pH 8.0, 150 mM NaCl, 2.0 mM CaCl) were prepared, onto which an equal volume of erythrocytes at a density of 5 × 10^8^ cells/ml were added, and the mixtures incubated at 37°C for 180 min under constant stirring. At the end of the incubation time, the plates were centrifuged (2000 x g, 10 minutes, 4 °C) and the supernatant scattering was measured at 700 nm. Alternatively, time course experiments were performed recording continuously the scattering signal at 700 nm. The blank (0% hemolysis) corresponded to erythrocytes incubated in buffer without toxin and 100%, and 100% hemolysis was obtained by adding Triton X-100 (0.1%) to the erythrocyte suspension.

### Cell culture

J774A.1 macrophages (ATTC, number TIB-67) were grown at 37°C in DMEM (Sigma Aldrich, USA) containing 10% (v/v) heat inactivated FBS (Thermo Fisher Scientific, USA), 6 mM L-glutamine (Thermo Fisher Scientific, USA), 0.2 % (v/v) MycoZapTM Prophylactic (Lonza, Switzerland) and Penicillin-Streptomycin (Sigma Aldrich, USA) (100 U/ml and 100 μg/ml respectively) in a 90% humidified atmosphere with 5% CO_2_.

### Measurement of cAMP

cAMP produced in cells was measured upon incubation of different ACT concentrations (25-200 ng/ml) with J774A.1 cells (5 × 10^5^ cells/ ml) for 30 min at 37°C. cAMP production was calculated by the direct cAMP ELISA kit (Enzo Lifesciences, USA).

### Measurement of ACT or mutant toxins binding to lipid membranes determined by flotation assays

Membrane association of ACT or ACT variants was assayed by flotation assay using large unilamellar vesicles (LUVs). LUVs were prepared following the extrusion method of Hope et al. [41]. Phospholipids and cholesterol were mixed in chloroform and dried under a N_2_ stream. Traces of organic solvent were removed by 2h vacuum pumping. Subsequently, the dried lipid films were dispersed in buffer and subjected to 10 freeze-thaw cycles prior to extrusion 10 times through 2 stacked polycarbonate membranes with a nominal pore size of 100 nm (Nuclepore, Inc., USA). Phospholipid concentration of liposome suspensions was determined by phosphate analysis [43].Liposome size was determined by Dynamic Light Scattering in Zetasizer Nano ZS (Malvern Panalytical Ltd, UK). Vesicle flotation experiments in sucrose gradients were subsequently performed following the method described by Yethon et al. [44]. In brief, 750 nM ACT and 1.5 mM LUVs (DOPC and DOPC.Chol 3:1 molar ratio, with 0.5% Rhodamine) are incubated for 30 minutes at 37°C, under stirring. 125 μl of each sample was adjusted to a sucrose concentration of 1.4 M in a final volume of 300 μl and subsequently overlaid with 400μl and 300μl layers of 0.8 and 0.5 M sucrose, respectively. The gradient was centrifuged at 436,000 g for 180 min in a TLA 120.2 rotor (Beckman Coulter, USA). After centrifugation, four 250 μl fractions were collected as depicted in **Fig. 1A**. The material adhered to the tubes was collected into a fifth fraction by washing with 250 μl of hot (100 °C) 1% (w/v) SDS. The different fractions were run on SDS-PAGE, and the presence of ACT was probed by Coomassie. Liposomes were monitored by measuring rhodamine fluorescence. The values displayed on the right correspond to the percentages of protein found co-floating with vesicles, calculated by densitometry. Densitometry of the bands was performed by using ImageJ software, and the percentage of binding to vesicles was calculated from the band intensities measured in the vesicle-floating fractions, relative to the sum of the intensities measured in all fractions. The results displayed are representative of at least two replicates.

## Author Contribution

JA and HO planned the experiments; JA and RA performed experiments and analysed the data; HO wrote the paper with contributions from all the authors.

## Acknowledgements

This study was supported by grants from the Spanish Ministerio de Economía y Competitividad BFU2017-82758-P (H.O.) and of Basque Government (Grupos Consolidados IT1264-19). JA was recipient of a fellowship from the University of Basque Country (UPV/EHU). RA holds a contract funded by the Fundación Biofisika Bizkaia.

